# No Trade-Offs Required: Cross-Feeding From Survival Alone

**DOI:** 10.64898/2026.08.07.743543

**Authors:** Samuel Rosean, Aviv Bergman

## Abstract

Cross-feeding relationships shape the composition of many microbial communities, yet the evolutionary processes that give rise to them remain poorly understood. Most theoretical and experimental work has therefore focused on minimal scenarios, particularly the stable cross-feeding polymorphisms that evolve in asexual populations growing on a single energy source (Helling et al., 1987). Yet replicate experiments do not always produce cross-feeding populations, raising the question of why genetically identical populations evolving under identical conditions can follow different evolutionary trajectories (Treves et al., 1998). Here we present a bare-bones agent-based model of evolution in a chemostat. We show that selection for energy acquisition alone is sufficient to promote the evolution of cross-feeding, without invoking mechanisms specific to metabolic exchange. The resulting communities nevertheless differ across replicate simulations, reproducing the qualitative variability observed experimentally.

**Significance Statement:** Microbial communities often depend on cross-feeding, in which one cell’s metabolic product becomes another’s energy source. Existing explanations typically invoke trade-offs between metabolic tasks or other mechanisms specific to cross-feeding itself. Using large-scale in silico simulations of evolution in a chemostat, we show that no such explanation is required. A population that competes for metabolic energy by utilizing a primary resource and then releasing a product that may itself serve as an energy source can evolve into a mixed population of organisms that specialize in the primary resource alongside others that specialize in the secondary one. Energy-based probabilistic death and reproduction are sufficient to produce this coexistence and to reproduce the mixed outcomes seen in laboratory evolution experiments.

## Introduction

Cross-feeding relationships, in which one population’s metabolic by-product becomes a resource for another, shape the composition of many microbial communities [1]. Much of what is known about their evolutionary origin comes from long-term experiments in the model organism *Escherichia coli* [2, 3]. Until the mid-20th century, the classical view held that asexual microbial populations would remain genetically uniform—effectively a single clone between successive adaptive replacements [4]. From the 1950s onward, serial transfer experiments in *E. coli* suggested that when new variants did arise, evolving asexual populations were nevertheless dominated by successive clonal replacements, each adaptive sweep fixing one lineage and purging accumulated mutants [5]. With the emergence of neutral evolutionary theory, it was supposed that many variants remained in equilibrium until the environment or a specific variant conferred a fitness difference [6, 7]. By the early 1980s, experimental evidence began to indicate that in the model asexual species *E. coli*, there was more genetic diversity among strains than in most eukaryotic species [7]. Starting in 1987, experiments spanning hundreds of generations in glucose-limited chemostats and long-term serial batch cultures have shown that *E. coli* populations could develop stable polymorphisms and genotypic diversification [2, 8]. It was sometimes observed that some subgroups excreted a metabolite that others consumed, producing cross-feeding between variants [2, 8, 9]. These observations ran against the Competitive Exclusion Principle [10, 11], which holds that two populations cannot sta-bly coexist on a single resource. In *E. coli*, stable polymorphism was thought to arise from differences in growth rate on the primary resource, glucose, and in excretion of an intermediate product, generally acetate or glycerol [8, 12–14]. This is an example of what Glen D’Souza et al. call unidirectional by-product cross-feeding [1], wherein one population releases a metabolic by-product that confers no benefit on the producer but benefits another—ecologically, a commensalism. This resolves the exclusion principle’s tension by carving two distinct niches out of a single inflowing resource.

Confoundingly, however, cross-feeding relationships did not always emerge in long-term evolution experiments under the same conditions, as in the 1998 experiment by Treves et al., which found that only 6 out of 12 identical populations of *E. coli* developed cross-feeding relationships after 1,750 generations [12].

Does this imply that replicates diverge due to sensitivity to initial conditions or because of a historical contingency in the sequence of vital events [15–17]? If a cross-feeding-specific trade-off alone governed emergence, replicate lines would reliably produce crossfeeding under matched conditions; we instead ask whether death and reproduction based on metabolic output alone can produce this mixture of outcomes.

Most work on this topic agrees on a minimal setup: organisms with two linked metabolic tasks (Figure 1), in which Task A and/or Task B can supply energy [8, 18, 19]. We refer to this general system as the *Minimal Chemostat Cross-Feeding Model* (MCCM). What exactly each part of this system represents and how it should be modeled—say, whether Task A represents a single action or a series of linked chemical reactions—varies widely among approaches to modeling this problem [18–22], but the core description remains the same.

**Figure 1:**
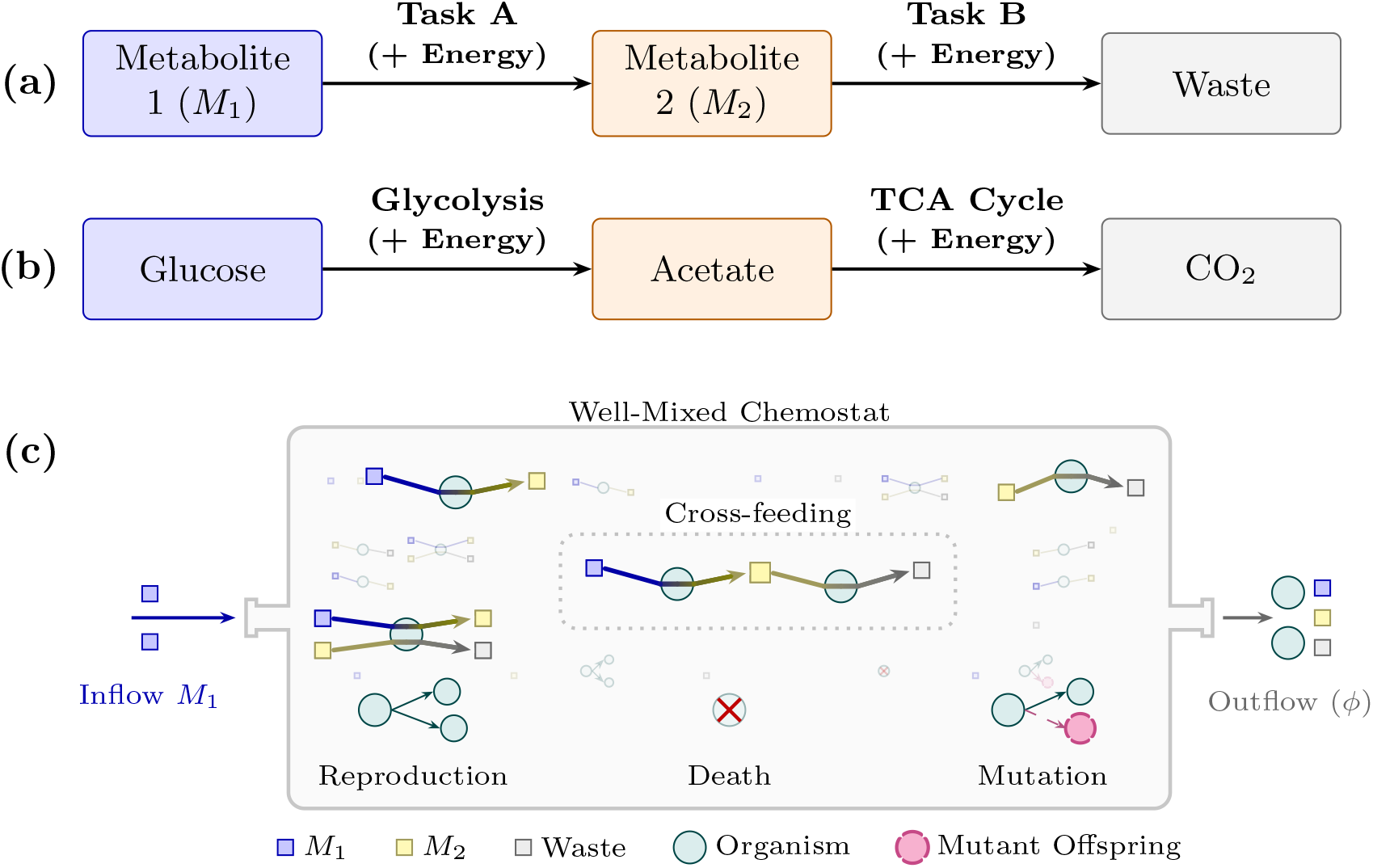
Schematic of the MCCM metabolic motif, biological interpretation, and well-mixed chemostat setup. **a)** Task A converts *M*_1_ to *M*_2_; Task B converts *M*_2_ to waste; each task yields energy for the organism. **b)** Simplified *E. coli* reading: Glucose → Acetate → CO_2_. **c)** *M*_1_ (blue) is fed into the chemostat; organisms convert inflowing *M*_1_ to *M*_2_ (yellow), *M*_2_ to waste (grey), or a combination of both. In the center, a cross-feeding relationship is emphasized,with the product of one organism’s metabolism being consumed by another. Illustrative reproduction, death, and trait mutation events are shown along the lower chemostat; outflow (*ϕ*) removes metabolites and organisms at a constant rate.

Previous work has often treated mechanisms intrinsic to the cross-feeding interaction as the main drivers of its emergence. Doebeli’s trade-off curve is a leading example, in which committing more to one task non-linearly worsens the capacity of the organism to perform the other, which can favor diversification even in a homogeneous environment. In fact, Doebeli found that such non-linear trade-offs between tasks were necessary for cross-feeding to arise in his models [18, 23, 24]. This trade-off-based account now forms a widely cited framework for thinking about the evolutionary emergence of cross-feeding and its dynamics [25, 26]. Cross-feeding-related mechanisms like trade-offs [18, 23, 26], proteome allocation [22], and the diffusion of metabolites [27] likely play a role in emergence, but it remains unclear whether they are sufficient to explain experimental variability [12], or even strictly necessary for cross-feeding to arise in general.

Theoretical minimal chemostat cross-feeding models have likewise focused on describing regimes where cross-feeding can and cannot emerge, rather than the specific dynamics of that evolution. The predominant method [18, 19] has been invasion analysis—including pairwise invasion or invasibility plots—wherein the fitness (or, in chemostat formulations, growth rate) of a resident genotype is compared with that of a rare invader; if the invader has a higher growth rate, it may displace the resident, or a new population may be built with the invader balanced with the current resident. While this approach has proven useful in identifying possible environmental and internal drivers of cross-feeding, it assesses a rare invader against a resident assumed to sit at equilibrium, and so characterizes whether cross-feeding can invade an established community rather than how it emerges while the population is still far from that equilibrium. It also ignores the fact that asexual populations are not genotypically homogeneous, and that multiple competing variants likely coexist in the population.

Here, rather than characterizing which trait compositions are stable once reached, we instead simulate the full generation-by-generation development of an initially homogeneous population using an agent-based model. We then measure the cross-feeding relationships in that final generation, scoring emergence rather than asymptotic stability. This allows us to ask whether probabilistic differential death and reproduction based on each organism’s energy is enough, on its own, to give rise to a population of primary- and secondary-resource users, and to achieve the non-deterministic, only partially repeatable character observed across Treves et al.’s replicate lines. This approach allows us to consider the generation-by-generation costs of spending energy on reproduction and bearing the risks of death, complexities that allow for non-ideal trajectories and local optima that are underconsidered in previous work.

## Results

Consider a population in which each organism performs two linked metabolic tasks (Figure 1): Task A converts the inflowing resource *M*_1_ into *M*_2_, and Task B converts *M*_2_ into waste, with each task yielding energy for the organism that performs it. Commitment to the two tasks is captured by a pair of traits, *A*_*i*_ and *B*_*i*_ = 1 − *A*_*i*_, so that organism *i* may allocate effort to Task A, Task B, or any combination of the two; since *B*_*i*_ is fully determined by *A*_*i*_, this reduces each organism to a single genotype value, *A*_*i*_ ∈ [0, 1]. Each simulation begins as a homogeneous population with *A*_*i*_ = *A*_0_ for all *i*, where *A*_0_ is sampled from the range given in Supplementary Table S1. From this starting point, we ask under what conditions the population diversifies: whether one portion comes to rely primarily on Task A and *M*_1_, while another relies primarily on Task B and *M*_2_.

Each generation, an organism stores the energy earned from Task A and/or Task B minus a fixed per-generation maintenance cost. Death and reproduction for each organism are then determined by independent random draws, with probabilities that depend on the energy remaining after this update. When reproduction occurs, the offspring’s genotype may mutate away from the parent’s, with probability *µ*_m_ and step size *σ*_m_, both sampled from Supplementary Table S1. We then ask whether changing how these energy-dependent death and reproduction probabilities are defined affects the frequency with which cross-feeding outcomes emerge from an initially homogeneous population.

### Death, reproduction, and chemostat outflow

Traits *A*_*i*_ and *B*_*i*_ set each organism’s relative demand for Tasks A and B, whereas *A*_*i*_ and ℬ_*i*_ are the units of metabolism it actually completes in a generation. Because metabolites are drawn from shared pools in the well-mixed chemostat, total demand can exceed available *M*_1_ or *M*_2_. In this case, each organism receives a share of the relevant pool proportional to its commitment to that task. An organism highly invested in Task A will receive a larger share of the *M*_1_ pool than an organism more invested in Task B, reflecting its greater commitment to Task A and thus a larger proportional draw on that pool:

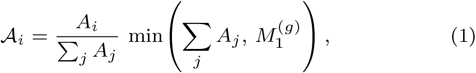

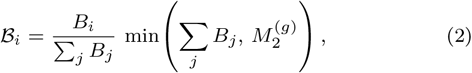

where 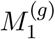 is environmental *M*_1_ after inflow, and 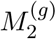 is the shared *M*_2_ pool when Task B runs, including *M*_2_ released by Task A that generation. Competition is pool-specific: Task A users compete for *M*_1_ (Eq. (1)), Task B users for *M*_2_ (Eq. (2)), so dominating *M*_1_ does not remove the cross-feeding niche on *M*_2_. Let 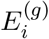 be organism *i*’s energy after metabolism and maintenance in generation *g*:

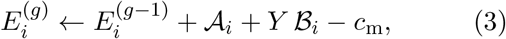

where *c*_m_ is the per-generation maintenance cost and *Y* is the relative ATP yield of Task B versus Task A in Eq. (3)—not the free-energy content of *M*_2_ relative to *M*_1_. For the glucose→acetate motif in Figure 1b, Task A is glycolysis to acetate and Task B is oxidation of that acetate through the TCA cycle. Theoretical yields make Task B the higher-ATP arm—a few ATP from producing acetate versus several-fold more from oxidizing it [28, 29]—so *Y* ∼ 2–4 is a reasonable range for this motif. Death and reproduction are independent Bernoulli draws with probabilities *P*_death,*i*_(*E*_*i*_) and *P*_repro,*i*_(*E*_*i*_). Each survivor is then removed by outflow with probability *P*_outflow,*i*_ = *ϕ*. In the neutral baseline, *P*_death,*i*_ = 0, so removal comes only from outflow (Supplementary Information, Neutral baseline).

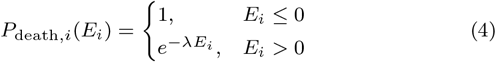

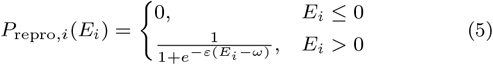

We compare a neutral baseline with a combined energy-dependent death-and-reproduction regime (Figure 2). In **Neutral Regime**, there is no energydependent death, and reproduction is set to balance chemostat outflow, so removal and reproduction do not select on metabolic success. In **Selection Regime**, low-energy organisms die preferentially (Eq. (4)) and high-energy organisms reproduce preferentially (Eq. (5)).

**Figure 2:**
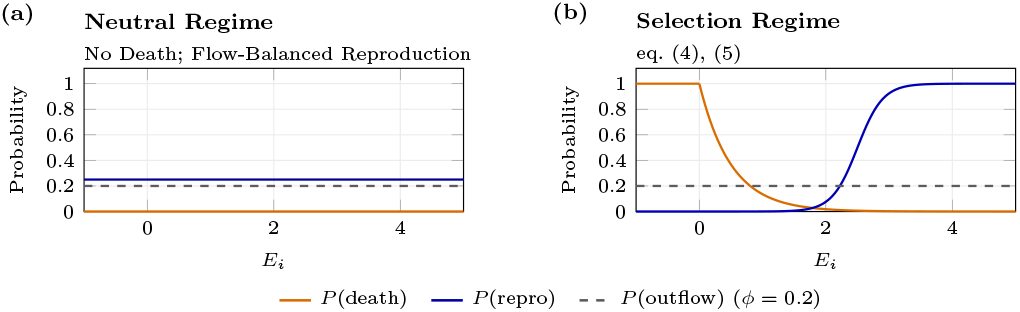
Death-and-reproduction configurations plotted as *P*_death,*i*_(*E*_*i*_), *P*_repro,*i*_(*E*_*i*_), and *P*_outflow,*i*_ versus post-metabolism energy *E*_*i*_. **a) Neutral Regime:** No Death with Flow-Balanced Reproduction. **b) Selection Regime:** Eq. (4) and Eq. (5) (*λ*=2, *ε*=5, *ω*=2.5).

### Simulation campaigns and cross-feeding criterion

In each simulation, following Supplementary Figure S1, an initial population of *N*_0_=100 homogeneous organisms evolves in a chemostat for *G*=1000 generations under one of the configurations in Figure 2 and a fixed task energy yield value *Y* ∈ {10^*−*4^, 0.1, 0.25, 0.5, 0.75, 1, 3, 5, 7, 10}. Each simula-tion has an independently sampled parameter vector drawn from Supplementary Table S1. Running 1000 of these simulations forms a batch, and we then run *B*_c_=100 independent batches. After the final generation, each run is scored with three tests: (i) whether the population persists, (ii) whether organisms have specialized into different metabolic roles, and (iii) whether secondary-resource use reflects genuine cross-feeding. All tests use each organism’s realized throughputs 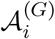 and 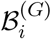 at generation *G* (Eq. (1) and Eq. (2) with *g*=*G*). The batch hit count, *H*_*b*_, is the number of simulations that passed all three tests (notation summarized in Supplementary Table S2).

### *Persistence* Is there still a population to score?

We require *N*_*G*_ *>* 50, where *N*_*G*_ is the number of organisms remaining at generation *G*, so we score only populations large enough that an apparent exchange fraction is unlikely to be driven by small-population drift alone.

### *Specialization*. Have organisms diverged in metabolic role?

For each organism, the Task B share of its activity is

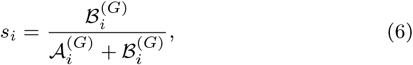

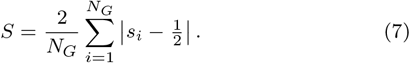

*S*=0 means that every organism is balanced (*s*_*i*_=0.5); *S*=1 means that every organism is committed to one task alone. We require *S >* 0.3 (Eq. (7)), which means a population that is not entirely task-balanced, which excludes homogeneous populations before the more expensive *Departure from Neutral Drift* test.

### *Departure from Neutral Drift*Is there more secondary-resource cross-feeding than would be expected from neutral genetic drift alone?

We first measure how much of the population’s Task B activity in the final generation depends on *M*_2_ produced by other organisms rather than by each organism itself. An organism with 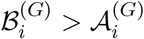 completes more Task B (consuming *M*_2_) than Task A (producing *M*_2_), so the surplus it uses must have been released by others. For each such organism, we define the Task B excess

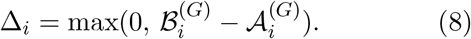

Organisms that do not consume more *M*_2_ than they produce contribute zero. The total amount of Task B excess in the population is Δ^tot^ = Σ _*i*_ Δ_*i*_. The total Task B throughput, regardless of whether it is in excess, is 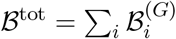. The exchange fraction

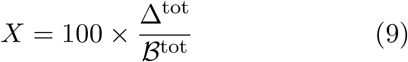

is then the percentage of all Task B throughput that is accounted for by that excess—equivalently, the share of Task B activity that depends on cross-fed *M*_2_. A high *X* therefore means that much of the population’s use of *M*_2_ depends on metabolites produced by others, not on self-produced *M*_2_.

A high *X* alone is not evidence of adaptive cross-feeding, since neutral demographic drift, in the absence of energy-dependent selection, can also produce a nonzero exchange fraction. We therefore compare each selection run against a matched set of *R*=100 neutral drift runs (Supplementary Information, *Neutral drift runs*) that replay the same history of event counts under no selection, yielding a null distribution of exchange fractions *X*^(*j*)^. The neutral-drift percentile

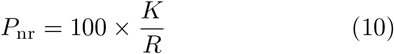

gives the fraction *K* of those *R* neutral drift runs with *X*^(*j*)^ *< X*; we treat the observed exchange fraction as significant when *P*_nr_ *>* 95%.

### Fixed-*Y* campaign outcomes and re-screens

Figure 3a demonstrates a relationship between combined energy-dependent death and reproduction and cross-feeding outcomes. **Neutral Regime** remains near ≈ 12.5 mean batch hits (≈ 1.3% per simulation) across the fixed-*Y* grid. At the lowest yield (*Y* =10^*−*4^), **Selection Regime** does not exceed this baseline, but from *Y* =0.1 upward it does, and the separation from **Neutral Regime** generally increases with *Y* .

**Figure 3:**
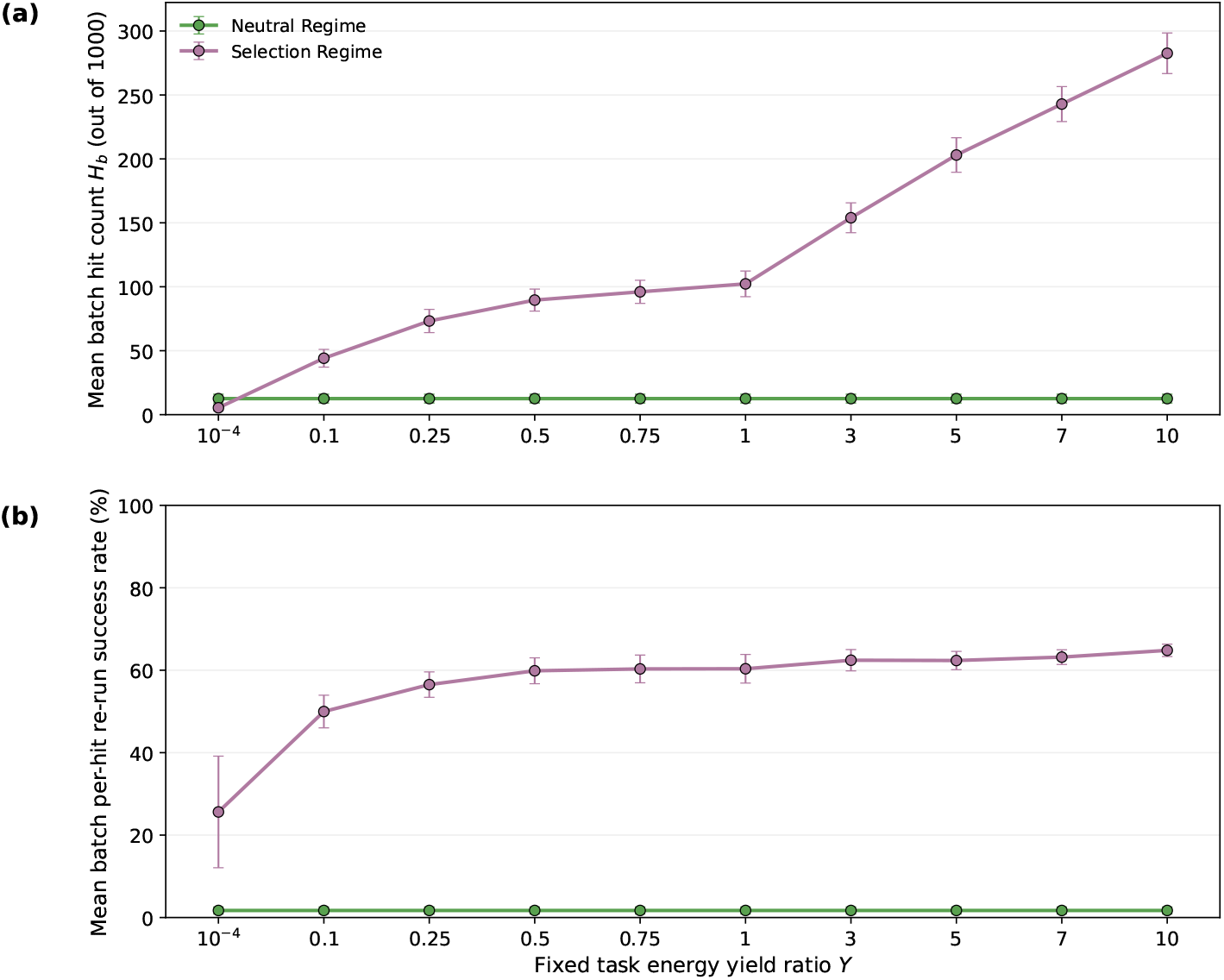
Fixed-*Y* campaign outcomes and hit re-screen confirmation for **Neutral Regime** and **Selection Regime** (Figure 2; *Y* ∈ {10^−4^, 0.1, 0.25, 0.5, 0.75, 1, 3, 5, 7, 10}). (a) Mean batch hit counts, *H*_*b*_, across *B*_c_=100 independent batches per configuration (*N*_sim_=1000 simulations per batch); error bars, ± SD across batches. (b) Mean batch per-hit re-run success rate across the same campaigns; error bars, ± SD across the same *B*_c_=100 batches. Each primary-batch hit was re-run *N*_re_=20 times with an identical parameter vector but a different seed, and the per-hit rate is the fraction of those re-runs that pass the cross-feeding criterion again. Per-hit success-count distributions are in Supplementary Figure S2.

We then re-screened every primary-batch hit in these fixed-*Y* campaigns to ask whether the same parameter vector would produce cross-feeding again with a different random seed driving the dynamics. Each hit was re-run *N*_re_=20 times with all settings held fixed except the seed. Figure 3b plots the mean per-hit re-run success rate; per-hit success-count distributions across batches are in Supplementary Figure S2. **Selection Regime** hits re-confirm substantially more often than **Neutral Regime** hits at every *Y* .

For a generation-by-generation view of how such **Selection Regime** hits unfold, Supplementary Figure S3 shows example trajectories of mean trait *A*, population size, and the neutral-drift percentile *P*_nr_ for four randomly selected hit runs.

This pattern indicates that many **Selection Regime** cross-feeding outcomes are tied to the sampled parameters and the shape of the death and reproduction rate curves rather than to a single lucky draw. **Selection Regime** hits re-confirm at ≈ 26– 65% across the full *Y* grid (rising to ≈ 60–65% for *Y* ≥ 0.5) while **Neutral Regime** hits re-confirm at only ≈ 2%. Re-run success nevertheless remains well below 100% even in the most robust regimes, so parameter sets can be partially reproducible while outcomes are still not fully deterministic across seeds. This demonstrates that selection factors and mechanics not directly related to cross-feeding can increase both the likelihood of cross-feeding emergence and parameter-linked robustness at particular points in parameter space, yet these probabilistic generational dynamics still allow non-deterministic outcomes.

## Discussion

Prior MCCM work uses invasion analysis to ask which genotype compositions are stable once reached [18, 19, 22], not whether evolution actually reaches them from a homogeneous start. Our simulations include a cost that those comparisons omit: at division the parent’s stored energy is split equally between parent and offspring—a standard conservation rule when pole age is not tracked—so reproduction carries an immediate energy cost. Real cells can also partition damage and storage asymmetrically, and death can depend on age as well as energy [30]; we omit those asymmetries. Because of that generational accounting, a one-step growth advantage need not persist, and trajectories need not maximize fitness at every step.

The experiment by Treves et al., which showed that identical chemostat lines differentially developed cross-feeding [12], raises a basic question about its origin: are these effects chaotic, i.e., dependent on imperceptibly minute differences in initial conditions between runs that are amplified over time, or are they stochastic and historical, depending on the actual sequence of vital-rate events that occur over evolutionary time? Our re-screen results speak directly to this distinction. **Selection Regime** hits re-confirmed at ≈26–65% across independent seeds under an otherwise identical parameter vector—far above the re-confirmation rate of **Neutral Regime** hits, yet still well below 100%. A purely chaotic account, in which outcome is fixed by initial conditions alone, would not predict this partial, parameter-linked reproducibility; a purely deterministic account would not predict its shortfall from 100%. The pattern instead indicates that the specific sequence of stochastic death and reproduction events realized over the course of a run—evolutionary contingency—is itself a source of variability that also fosters cross-feeding outcomes, alongside the sampled parameters. Previous models have identified conditions under which evolutionary branching and cross-feeding coexistence are expected [18, 19, 22], but because they do not address the temporal sequence or vital dynamics by which replicate populations traverse those conditions, they offer no account of the experimental variability Treves et al. observed. The possible mechanistic origins of such probabilistic death and reproduction are numerous, but a few examples include the inherent stochasticity of *E. coli* replication, which makes cell division initiate not at a constant time, but at a variable one [31], and starvation death in *E. coli*, which is set by active ion homeostasis that maintains plasmolysis, so that removal is energetically conditioned rather than fixed [32].

Trade-offs, proteome costs, and spatial structure likely still matter for maintaining cross-feeding once it exists [1, 18, 19, 22, 33]. They are not required to produce it: energy-dependent death and reproduction alone can diversify a homogeneous population into primary- and secondary-resource users, with the same partial repeatability seen in replicate experiments. Which secondary metabolites can support that diversification still depends on chemistry we do not model here—membrane permeability, environmental half-life, and toxicity of *M*_2_ among them [1, 14, 22]. A fuller account would therefore join the emergence mechanisms emphasized here with existing stability theory for established cross-feeding [34]. As agent-based metabolic and community-scale models of microbial interactions begin to incorporate evolution [35–37], an important design choice follows directly from this distinction: whether death and reproduction are treated as fixed demographic background or as stochastic processes in their own right. Our results argue for the latter. Even when the metabolic coupling is reduced to its simplest possible form, with no trade-off, proteome cost, or spatial structure, probabilistic vital dynamics alone shift the frequency and repeatability of cooperative outcomes. Models that extend flux-balance, spatial, or interaction-based community frameworks with evolutionary dynamics should therefore treat vital-rate stochasticity and generational history as first-class components of the evolutionary model, not as background noise or a secondary component of cross-feeding dynamics.

Read this way, the incomplete repeatability we observe is not a defect of the model but the expected signature of historical contingency: the same population, under the same survival pressures, arrives at cross-feeding or not depending on the particular sequence of reproduction and death events it happens to realize. Just as replaying evolution from a shared ancestor yields a metabolic innovation in some replicate *E. coli* lineages but not others [15, 16], cross-feeding here is reachable but not guaranteed—an emergent property of the probabilistic energetics of survival rather than of any mechanism dedicated to metabolic exchange.

Finally, while theoretical refinement is important, advancing first-principles accounts of cross-feeding emergence like the MCCM requires experimental paradigms that can produce robust enough evolutionary data to test these accounts. Extant data on the evolution of cross-feeding behaviors, and on how the relative abundance of cross-feeding relationships scales with a cross-fed metabolite’s energy yield (which would allow more direct comparison to our model), remain insufficient to validate our predictions. Theoretical expositions like ours can open the door to such paradigms; work that bridges this gap would make that validation possible, but would also open the door for a better evaluation of minimal cross-feeding models more broadly.

## Materials and Methods

### Simulation model

Our simulation models a minimal well-mixed chemostat, a step-by-step flowchart and chemostat snapshots visualizing the model can be seen in Supplementary Figure S1. Inside an enclosed environment, the population receives an inflow of resources and experiences a continuous outflow each generation. Inflow adds a constant amount of metabolite *M*_1_ into the environmental pool, in this case 100 units per generation. After metabolism, death and reproduction are drawn from the energy-dependent rate laws in Figure 2; chemostat outflow then removes surviving organisms with probability *ϕ* and scales environmental *M*_1_ and *M*_2_ by (1 − *ϕ*). The well-mixed environment gives all organisms equal access to environmental metabolites. Task traits and realized throughputs follow Eq. (1) and Eq. (2), and energy gain and maintenance follow Eq. (3), as described under *Death, reproduction, and chemostat outflow*. When an organism reproduces, its post-metabolism energy store is split equally between parent and offspring. In **Neutral Regime**, *P*_death,*i*_ = 0 and constant energy-independant reproduction is flow-balanced as *P*_repro,*i*_ = *ϕ/*(1 − *ϕ*), so expected population size is unchanged under chemostat outflow alone.

### Trait mutation upon reproduction

When an organism reproduces, parent and offspring each receive half of the parent’s stored energy, the offspring inherits the parent’s *A*_*i*_, and the offspring may then mutate independently with probability *µ*_m_ (Supplementary Table S1).

A naive mutation kernel—draw a uniform step of ±*σ*_m_ around the parental trait and then clip the result to [0, 1]—would introduce a spurious bias unrelated to selection. Near either boundary, clipping truncates outward moves but still allows inward ones, so accepted mutations are pushed toward the poles (*A*_*i*_ ≈ 0 or *A*_*i*_ ≈ 1). We instead draw the offspring trait directly from the *truncated uniform* window that remains feasible on [0, 1]:

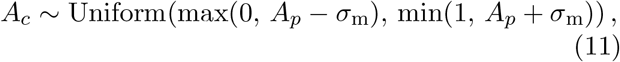

where *A*_*p*_ is the parental Task A trait, *A*_*c*_ is the offspring value, and *σ*_m_ is the sampled mutation scale. Because the draw is uniform over the clipped interval rather than over ±*σ*_m_ followed by clipping, boundary-near parents place slightly more probability on inward moves. That gives a mild inward pressure toward the trait interior and away from extreme specialization—that is, toward generalist, non-cross-feeding genotypes—without any explicit fitness term. Supplementary Information, *Neutral drift runs* applies the same kernel during reproduction steps.

### Neutral drift runs

For each **Selection Regime** run, we construct a set of neutral drift runs (Supplementary Information, *Neutral drift runs*) to estimate the null distribution of the exchange fraction under demographic drift alone. Each neutral drift run (i) starts from the same initial population as the selection run; (ii) reapplies, generation by generation, the same *counts* of deaths, reproduction events, outflow removals, and accepted mutations that occurred in the selection run, but reassigns *which* organisms are affected uniformly at random, holding the number of events fixed while reshuffling who is affected; and (iii)assigns Task A and Task B throughputs from the selection run’s final metabolite pools, with no energy-dependent death, reproduction, or metabolism. Neutral drift run *j* thus yields its own exchange fraction *X*^(*j*)^, the value that same history of event counts would produce under no selection. We draw a fixed number *R*=100 of neutral drift runs per selection run and let *K* denote the number with *X*^(*j*)^ *< X*, from which *P*_nr_ is computed via Eq. (10) (for additional details, see Supplementary Information, *Neutral drift runs*).

### Parameter sampling

Every simulation draws each varied parameter independently and uniformly from a given interval (Supplementary Table S1). Parameter draws, within-run stochastic dynamics, neutral drift runs, and re-screen trials use deterministic integer seeds (Supplementary Information, Deterministic Monte Carlo seeds). Death and reproduction follow one of the configurations in Figure 2; which rate laws are active and which of *λ, ε*, and *ω* are sampled are specified in Supplementary Table S1.

### Generative AI

During preparation of this work, the authors used Cursor (Anysphere, Inc.), an AI-assisted coding environment that routes prompts to large language models. Over 2025–2026 the models invoked through Cursor included Claude (Anthropic) and GPT (OpenAI) family models; the exact model identifier varied with Cursor platform updates and session settings and was not retained for every prompt. Cursor was used to assist with batch-processing, GUI, and analysis scripts along with manuscript editing and refinement. The primary simulation software was developed without AI assistance. All AI-suggested code and text were reviewed, tested, and edited by the authors, who take full responsibility for the published content.

## Data availability

Code and analysis scripts to reproduce the agent-based model and figures are available at https://github.com/AvivLab/Cross-Feeding-Evolution.

## Acknowledgments

We thank Libusha Kelly, Roger Chang, Sarah Wolfson, Grace Richmond, Kelsey Bledsoe, and Fitzgerald Small for support and discussion. We also thank William Chang and Gordon Huang. A.B. acknowledges the Albert Einstein College of Medicine, and the Harold and Muriel Block Foundation, for their generous support. We acknowledge generative AI use as described in Materials and Methods.

## Supporting Information

No Trade-Offs Required: Cross-Feeding From Survival Alone

This Supporting Information accompanies the main text. Simulation campaign structure and the cross-feeding criterion are described in the main-text Results (*Simulation campaigns and crossfeeding criterion*); fixed-*Y* hit rates and re-screens are in *Fixed-Y campaign outcomes and re-screens*. A hit refers to a simulation whose final population passes the three criterion of cross-feeding described in the main text. Rate laws and throughputs are introduced under *Death, reproduction, and chemostat outflow* ; trait mutation upon reproduction is described in Materials and Methods (*Trait mutation upon reproduction*). Extended methods below expand the scoring protocol (neutral baseline and neutral drift runs) and document the deterministic Monte Carlo seed formulas, followed by Supplementary Figures S1–S3 and Tables S1–S2 cited in the main manuscript. Throughout this SI we use the same configuration names as the main text: **Neutral Regime** and **Selection Regime**. For eas of interoperability equations reproduced from the main text retain their main-text numbers (Eq. 1, Eq. 2, …).

### Neutral baseline (flow-balanced reproduction)

Configurations combine energy-dependent death and reproduction rate laws with the auxiliary forms used in Figure 2 in the main text:

*Equations from main text* (Eqs. (4)–(5)).

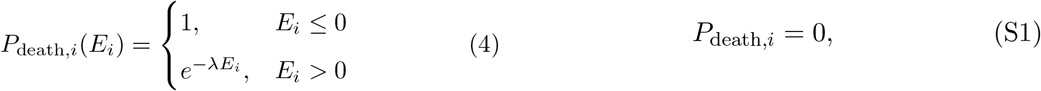

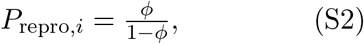

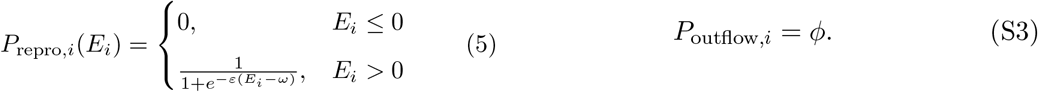

Configuration **Neutral Regime** sets *P*_death,*i*_ = 0 (SI Eq. S1), so per-organism removal in this baseline comes only from chemostat outflow (SI Eq. S3), not from an additional death draw. A flat, energy-independent death probability would act like extra dilution on top of outflow and is therefore omitted.

Each generation, after metabolism, surviving organisms first reproduce and are then subject to chemostat outflow (Fig. S1). On average, reproduction enlarges the population by a factor (1 + *P*_repro_), where *P*_repro_ is the per-organism energy-independent reproduction probability (SI Eq. S2 in **Neutral Regime**), and outflow then retains only a fraction (1 − *ϕ*) of that enlarged pool. The expected population size therefore updates as

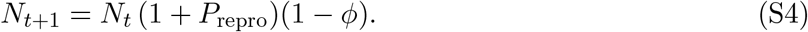

To keep expected population size steady under flow alone, we require (1 + *P*_repro_)(1 − *ϕ*) = 1, which gives *P*_repro_ = *ϕ/*(1 − *ϕ*). In sweeps, *ϕ* is sampled uniformly on [0, 0.5] (Table S1), so the corresponding flow-balanced reproduction rate *P*_repro_ = *ϕ/*(1 − *ϕ*) ranges from 0 to 1. Since this correction targets only the average population change, it does not guarantee stability in any individual run; neutral populations can therefore collapse or grow.

### Neutral drift runs

Neutral drift runs, which are distinct from the neutral regime, are created to replicate a specific run’s vital-events excluding energetics, metabolism, and selection. The neutral-drift runs generate the null distribution against which the vital runs are compared, providing the basis for the bootstrap procedure used to assess the statistical significance of the vital-run results. During each selection run, the model records a per-generation *change history*. For generation *g* = 1, …, *G*, let *D*_*g*_ denote the number of deaths, *U*_*g*_ the number of reproduction events, *F*_*g*_ the number of chemostat-flow removals, and *M*_*g*_ the number of accepted trait mutations (offspring whose *A*_*i*_ actually changed after reproduction).

Each neutral drift run proceeds as follows.

1. **Initialize Traits**. Start from the selection run’s homogeneous initial population, with *A*_*i*_ = *A*_0_ and *B*_*i*_ = 1 − *A*_0_ for all *i* (the same sampled *A*_0_ from the selection run as described in Table S1).
2. **Replay Demography and Mutations**. For each generation *g* in order, remove *D*_*g*_ organisms uniformly at random without replacement, add *U*_*g*_ offspring by drawing parents with replacement and copying their traits, apply trait mutations to exactly *M*_*g*_ of those offspring (chosen uniformly, with the same truncated-uniform kernel and *σ*_m_ as the simulator; see Eq. 11 in the main text), and remove *F*_*g*_ surviving organisms uniformly at random without replacement. This yields a final population with different trait values, but the same number of organisms as the selection run and an identical history of deaths, reproduction events, and flow removals.
3. **Neutral Pooled Task A and Task B Assignment**. Next we calculate throughputs for the neutral drift run. Because these runs omit energetics and metabolism, we use the selection run’s final environmental pool sizes after inflow (*M*_1_ and *M*_2_) and assign synthetic throughputs by the same homogeneous-pool rules as the simulator (see Eqs. 1–2 in the main text): Task A draws are allocated across organisms in proportion to *A*_*i*_; produced *M*_2_ is then returned to the shared pool, and Task B draws are allocated from that pool in proportion to *B*_*i*_. This produces per-organism throughputs 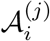 and 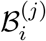.
4. **Exchange fraction**. Substitute 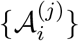 and 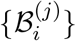 into Eqs. (8)–(9) to obtain *X*^(*j*)^.

Each neutral drift run *j* yields an exchange fraction *X*^(*j*)^. We draw a fixed number *R*=100 of neutral drift runs with deterministic seeds *j* = 0, 1, …, *R* − 1 (Deterministic Monte Carlo seeds). Let

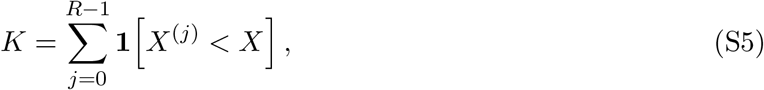

be the count where *X*^(*j*)^ *< X*. The neutral-drift percentile is

*Equation from main text (*Eq. (10)).

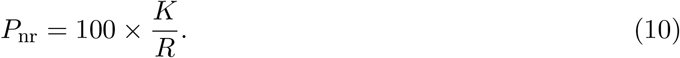

The selection run passes if *P*_nr_ *>* 95%.

### Deterministic Monte Carlo seeds

For completeness and to facilitate reproduction of our results, we describe here the exact procedure by which random seeds were chosen for the evolutionary dynamics. All reported campaigns use fixed integer seeds so that parameter draws, within-run stochastic dynamics, neutral drift runs, and re-screens are fully reproducible. Indices below are zero-based, matching the implementation: batches *b* = 0, …, *B*_c_ −1, simulations within a batch *r* = 0, …, *N*_sim_ −1, re-screen points *h* = 0, 1, …, and re-screen trials *t* = 0, …, *N*_re_ − 1.

Each primary-batch campaign is assigned a campaign base seed *s*_0_=1 (Table S1). Two fixed integer strides, *x* and *y* (Table S1), separate seeds across batches or re-screen parameter points (*x*) and across simulations or re-screen trials within a batch or point (*y*). To reproduce the reported seeds exactly, use *x*=10,007 and *y*=7,919. Batch *b* receives the batch seed

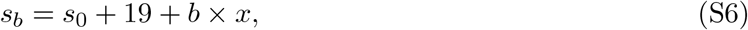

which seeds a NumPy pseudorandom generator used to draw that batch’s *N*_sim_ independent parameter vectors uniformly from the bounds in Table S1. Within batch *b*, simulation *r* is then assigned the dynamics seed

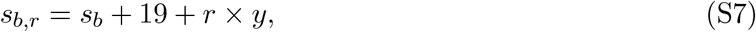

equivalently *s*_*b,r*_ = *s*_0_ + 38 + *b* × *x* + *r* × *y*. This value is passed to the simulator as the run’s random seed and governs all within-run stochastic events (Task A and Task B allocation, death, reproduction, outflow, and mutation). Substituting *s*_0_=1 gives *s*_*b*_ = 20 + *b* × *x* and *s*_*b,r*_ = 39 + *b* × *x* + *r* × *y*.

Because all primary-batch campaigns share the same base seed *s*_0_ and the task energy yield ratio *Y* does not affect **Neutral Regime** dynamics (*Y* enters only through energy gain and the energy-dependent rate laws that are absent in this configuration), the reported **Neutral Regime** campaigns at different fixed *Y* are identical: they use the same parameter draws and dynamics seeds and therefore yield *identical* results.

Neutral drift Monte Carlo draws (SI Eq. S5) use consecutive integer seeds *j* = 0, 1, …, *R* − 1 in order, with fixed *R*=100 as described under *Neutral drift runs*.

Hit re-screens hold the primary parameter vector fixed and vary only the dynamics seed. Rescreen trials use a separate base seed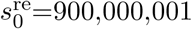. For unique parameter point *h* and trial *t*,

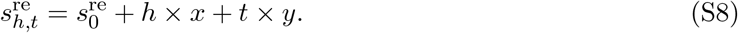

With *N*_re_=20 trials per point, *t* runs from 0 to 19.

**Table S1:** Model and sweep parameters for the reported Monte Carlo campaigns. Sampled rows apply only when the active configuration draws that quantity (see the configurations in Figure 2 in the main text). Additional symbols are in Table S2. Death and reproduction equations are in Eqs. (4)–(5) and SI Eqs. (S1)– (S3).

| Symbol | Parameter | Role | Range or value |
| --- | --- | --- | --- |
| *Death and reproduction* (Eqs. (4)–(5); SI Eqs. (S1)–(S3)) |  |  |  |
| $\lambda$ | Death decay | Controls $e^{-\lambda E_i}$ death above zero in **Selection Regime** (Eq. (4)) | Uniform(0.1, 20) |
| $\varepsilon$ | Repro. steepness | Sigmoid steepness in **Selection Regime** (Eq. (5)) | Uniform(0, 10) |
| $\omega$ | Repro. midpoint | Sigmoid midpoint energy in **Selection Regime** (Eq. (5)) | Uniform(0, 5) |
| *Evolution and flow* (sampled every simulation) |  |  |  |
| $c_m$ | Cost of life | Maintenance energy deducted each generation see Eq. 3 in the main text | Uniform(0.001, 0.05) |
| $\phi$ | Flow fraction | Chemostat outflow probability (SI Eq. S3); in **Neutral Regime**, also sets $P_{\text{repro},i}$ via SI Eq. S2 | Uniform(0, 0.5) |
| $\mu_m$ | Mutation rate | Probability that an offspring mutates upon reproduction | Uniform(0, 0.005) |
| $A_0$ | Initial Task A | Homogeneous starting genotype ($A_i=A_0$ at $g=1$) | Uniform(0, 1) |
| $E_{\text{init}}$ | Initial energy | Homogeneous starting energy for each organism at $g=1$ | Uniform(0, 20) |
| $\sigma_m$ | Mutation scale | Maximum trait step when a mutation is accepted | Uniform(0, 1) |
| *Fixed settings* |  |  |  |
| $Y$ | Task B/Task A energy yield ratio | Scales Task B energy yield relative to Task A in see Eq. 3 in the main text; one value per campaign, Figure 3 in the main text | $\{10^{-4}, 0.1, 0.25, 0.5, 0.75, 1, 3, 5, 7, 10\}$ |
| $N_0$ | Initial population | Organisms at $g=1$ | 100 |
| $G$ | Generations | Length of each run | 1,000 |
| $I_{M_1}$ | $M_1$ inflow | External glucose added per generation | 100 |
| *Cross-feeding criterion* (fixed thresholds; persistence, SI Eq. ??; specialization and Departure from Neutral Drift, Eqs. (6)–(10)) |  |  |  |
| — | Persistence | Minimum final population size $N_G$ | $> 50$ |
| — | Specialization | Minimum specialization index $S$ | $> 0.3$ |

**Table S1:** Continued from previous page.
| Symbol | Parameter | Role | Range or value |
| --- | --- | --- | --- |
| — | *Departure from Neutral Drift* | Minimum neutral-drift percentile $P_{nr}$ of the selection run's exchange fraction $X$ | $> 95$ |
| *Monte Carlo (fixed across sweeps)* |  |  |  |
| $N_{sim}$ | Simulations per batch | Independent parameter draws per batch | 1,000 |
| $B_c$ | Batches per campaign | Independent batches per configuration | 100 |
| $N_{re}$ | Re-screen trials | Fresh dynamics seeds per primary hit | 20 |
| $R$ | Neutral drift runs | Fixed number of null (neutral drift) runs per selection run<br>(*Neutral drift runs*) | 100 |
| $s_0$ | Campaign base seed | Root seed for primary-batch parameter draws and dynamics<br>(*Deterministic Monte Carlo seeds*) | 1 |
| $x$ | Batch/point seed stride | Stride across batches or re-screen parameter points (SI Eq. S6, SI Eq. S8) | 10,007 |
| $y$ | Simulation/trial seed stride | Stride across simulations within a batch or re-screen trials within a point (SI Eq. S7, SI Eq. S8) | 7,919 |
| $s_0^{re}$ | Re-screen base seed | Root seed for hit re-screen trials (*Deterministic Monte Carlo seeds*) | 900,000,001 |

**Table S2:** Notation for model state, outcome metrics, and neutral drift runs. The selection run is the evolutionary simulation being scored; neutral drift runs are its matched nulls. Sampled and fixed sweep parameters (*λ, ϕ, A*_0_, *N*_0_, *G, N*_sim_, *B*_c_, *x, y*, etc.) are in Table S1.

| Symbol | Quantity | Role |
| --- | --- | --- |
| *Indices and model state* |  |  |
| $i$ | Organism index | Labels per-organism quantities |
| $g$ | Generation index | Loop counter, $g = 1, \dots, G$; $g=1$ is run initialization |
| $M_1, M_2$ | Pool metabolites | Environmental substrates for Tasks A and B |
| $A_i, B_i$ | Task A and Task B traits | Task A and Task B commitment; $B_i=1-A_i$ |
| $\mathcal{A}_i, \mathcal{B}_i$ | Task A and Task B throughputs | Realized Task A and Task B units completed per generation |
| $\mathcal{A}_i^{(G)}, \mathcal{B}_i^{(G)}$ | Final throughputs | Throughputs at generation $G$ used in cross-feeding scoring |
| $E_i, E_i^{(g)}$ | Post-metabolism energy | Energy after metabolism and maintenance; vital-rate input |
| $P_{death,i}$ | Death probability | Per-organism probability of death each generation; constant or as a function of $E_i$ |
| $P_{repro,i}$ | Reproduction probability | Per-organism probability of producing one offspring each generation; constant or as a function of $E_i$ |
| $P_{outflow,i}$ | Outflow probability | Chemostat removal after death and reproduction |
| *Outcome metrics and cross-feeding criterion* |  |  |

**Table S2:** Continued from previous page.
| Symbol | Quantity | Role |
| --- | --- | --- |
| $b$ | Batch index | $b = 0, \dots, B_c - 1$ (zero-based; campaign summaries label batches $1, \dots, B_c$) |
| $H_b$ | Batch hit count | Simulations passing all three criteria (persistence, specialization, and *Departure from Neutral Drift*) in batch $b$ |
| $N_G$ | Final population size | Organisms remaining at generation $G$ (persistence uses $N_G > 50$) |
| $s_i$ | Task B share | $s_i = \mathcal{B}_i^{(G)} / (\mathcal{A}_i^{(G)} + \mathcal{B}_i^{(G)})$ |
| $S$ | Specialization index | Mean absolute deviation of $s_i$ from balanced (0.5); pass if $S > 0.3$ |
| $\Delta_i$ | Task B excess | $\max(0, \mathcal{B}_i^{(G)} - \mathcal{A}_i^{(G)})$ for organism $i$ (zero if not in excess) |
| $\Delta^{\text{tot}}$ | Total Task B excess | Total amount of Task B excess in the population, $\sum_i \Delta_i$ |
| $\mathcal{B}^{\text{tot}}$ | Total Task B throughput | Total Task B throughput, regardless of whether it is in excess, $\sum_i \mathcal{B}_i^{(G)}$ |
| $X$ | Exchange fraction | Percentage of all Task B throughput accounted for by that excess, $100\Delta^{\text{tot}}/\mathcal{B}^{\text{tot}}$ |
| $P_{\text{nr}}$ | Neutral-drift percentile | Percentage of neutral drift runs with exchange fraction below the selection run, $100K/R$; pass if $P_{\text{nr}} > 95$ |
| *Neutral drift runs (change history and Monte Carlo null)* |  |  |
| $D_g$ | Deaths | Organisms removed by death in generation $g$ (count fixed from the selection run) |
| $U_g$ | Reproduction events | Offspring added by reproduction in generation $g$ (count fixed from the selection run) |
| $F_g$ | Outflows | Organisms removed by chemostat outflow in generation $g$ (count fixed from the selection run) |
| $M_g$ | Mutations | Offspring whose $A_i$ changed after reproduction in generation $g$ (count fixed from the selection run) |
| $R$ | Neutral drift count | Fixed number of null runs used for $P_{\text{nr}}$ ($R=100$) |
| $K$ | Below-selection count | Number of null runs with exchange fraction $X^{(j)} < X$ |
| $j$ | Neutral drift seed index | Null-run index $j = 0, 1, \dots, R - 1$ |
| $X^{(j)}$ | Neutral drift exchange fraction | Exchange fraction from null run $j$ |
| $\mathcal{A}_i^{(j)}, \mathcal{B}_i^{(j)}$ | Neutral drift throughputs | Synthetic Task A and Task B throughputs from null run $j$ |
| *Deterministic Monte Carlo seeds (zero-based indices)* |  |  |
| $s_0$ | Campaign base seed | Root seed for primary campaigns ($s_0=1$) |
| $x$ | Batch/point seed stride | Stride across batches $b$ or re-screen points $h$ ($x=10,007$) |
| $y$ | Simulation/trial seed stride | Stride across simulations $r$ or re-screen trials $t$ ($y=7,919$) |
| $s_b$ | Batch seed | Parameter-draw RNG seed for batch $b$ |
| $r$ | Simulation index | $r = 0, \dots, N_{\text{sim}} - 1$ within batch $b$ |
| $s_{b,r}$ | Run dynamics seed | Within-run stochastic seed for simulation $r$ in batch $b$ |
| $s_0^{\text{re}}$ | Re-screen base seed | Root seed for re-screen trials ($s_0^{\text{re}}=900,000,001$) |
| $s_{h,t}^{\text{re}}$ | Re-screen trial seed | Dynamics seed for parameter point $h$, trial $t$ |

**Table S2:** Continued from previous page.
| Symbol | Quantity | Role |
| --- | --- | --- |
| $h$ | Re-screen point index | Unique primary parameter vector index ($h=0, 1, \dots$) |
| $t$ | Re-screen trial index | Fresh-seed trial within a re-screen ($t=0, \dots, N_{\text{re}} - 1$) |

**Figure S1:**
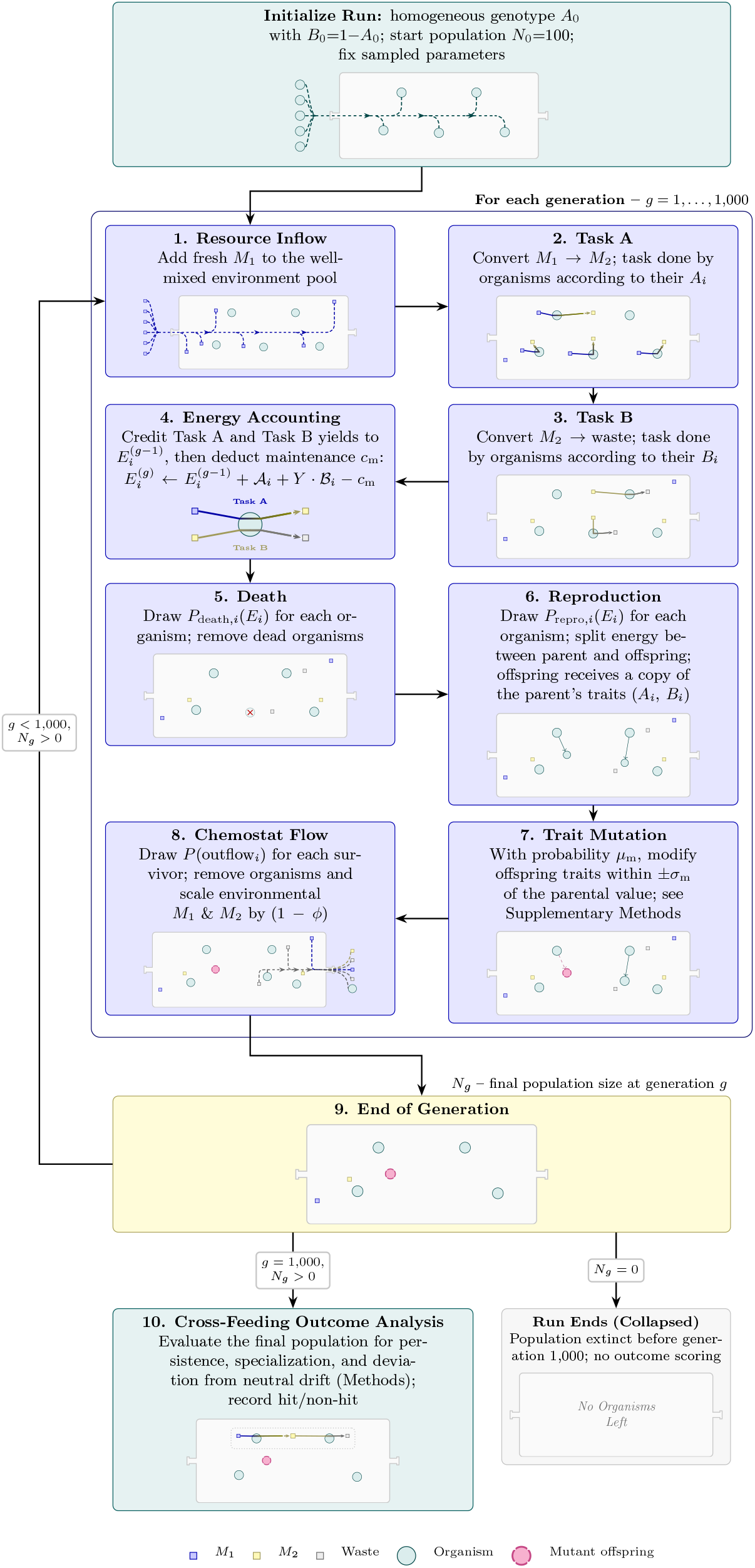
Agent-based simulation flow for each simulation. After initialization, each generation runs steps 1–8; step 7 summarizes trait mutation in general terms (details in Materials and Methods, *Trait mutation upon reproduction*). Step 9 tests for population collapse and whether *g*=1,000, looping back to step 1 when more generations remain or ending the run if the population has reached zero. After the final generation,step 10 scores the population against the cross-feeding criterion (main-text Results, *Simulation campaigns and cross-feeding criterion*). Each step box includes a chemostat snapshot below the written description; the boxed region covers the per-generation loop (steps 1–9).

**Figure S2:**
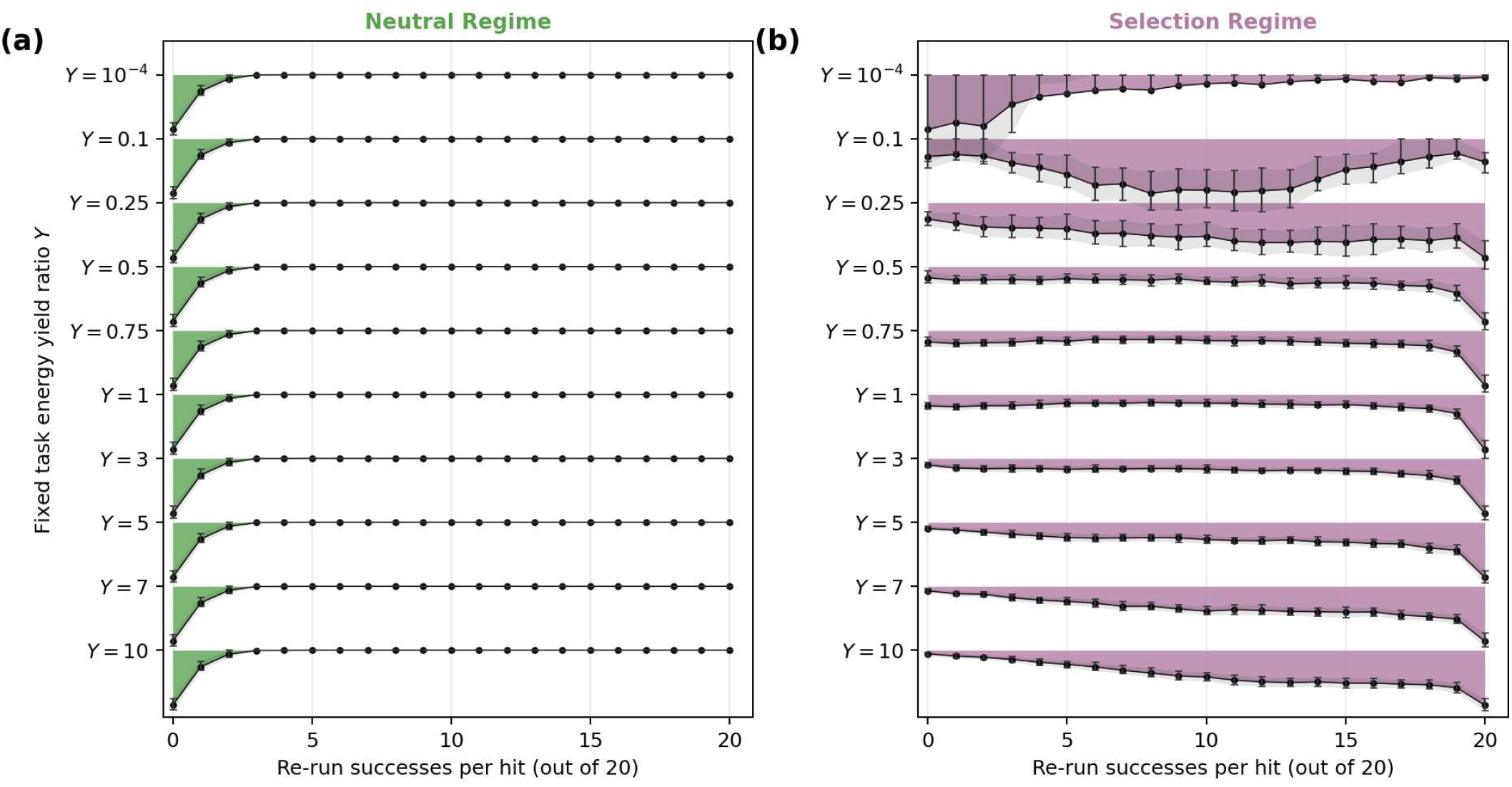
Per-hit re-run success-count distributions for the fixed-*Y* hit re-screens summarized in Figure 3b in the main text. Each primary-batch hit was re-run *N*_re_=20 times with all settings held fixed except the seed. For each success count *k* ∈ {0, …, *N*_re_}, ridges show the mean per-batch density at *k* (peak-normalized for visual comparison); vertical error bars, quartile range (Q1–Q3) across *B*_c_=100 independent batches per configuration. **a) Neutral Regime. b) Selection Regime**. Within each panel, ridges are ordered by fixed *Y*.

**Figure S3:**
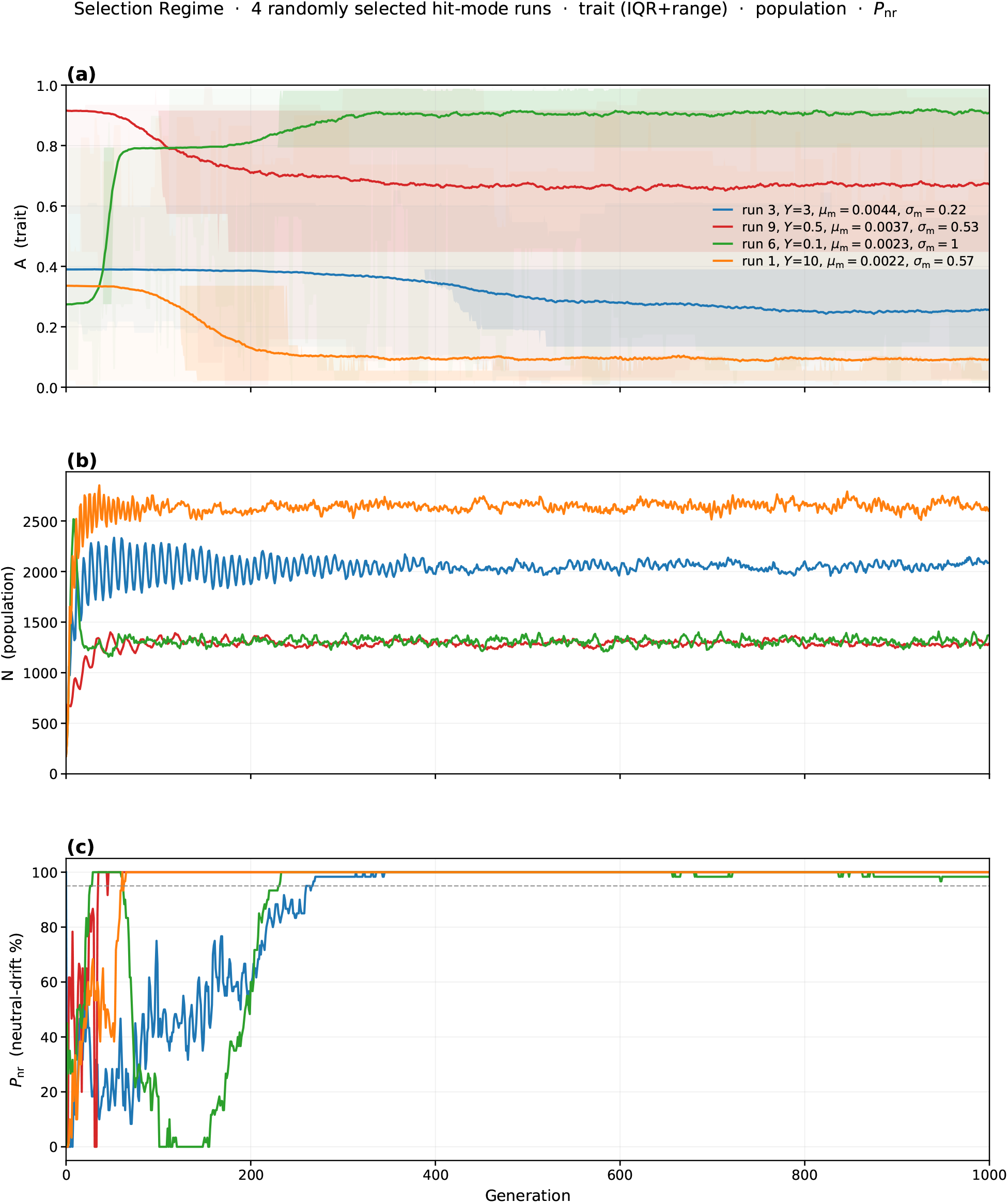
Generation-resolved trajectories for four randomly selected **Selection Regime** runs from the hit-mode trajectory sessions (*Y* ∈ {10^−4^, 0.1, 0.25, 0.5, 0.75, 1, 3, 5, 7, 10}; random seed 0). The figure stacks three time series. **a)** Mean Task A trait with IQR and full-range bands (commitment to primary-resource metabolism; *A*=1 is Task A only, *A*=0 is Task B only). **b)** Population size *N* . **c)** Neutral-drift percentile *P*_nr_ of the exchange fraction at that generation (percentage of contemporaneous neutral drift runs with a lower exchange fraction; dashed line at the 95% pass threshold). Legend entries identify each run index, fixed *Y*, mutation rate *µ*_m_, and mutation scale *σ*_m_.

## Notes

https://github.com/AvivLab/Cross-Feeding-Evolution

### Competing Interest Statement

The authors have declared no competing interest.

